# Allele frequency divergence reveals ubiquitous influence of positive selection in *Drosophila*

**DOI:** 10.1101/2021.03.15.435474

**Authors:** Jason Bertram

## Abstract

Resolving the role of natural selection is a basic objective of evolutionary biology. It is generally difficult to detect the influence of selection because ubiquitous non-selective stochastic change in allele frequencies (genetic drift) degrades evidence of selection. As a result, selection scans typically only identify genomic regions that have undergone episodes of intense selection. Yet it seems likely such episodes are the exception; the norm is more likely to involve subtle, concurrent selective changes at a large number of loci. We develop a new theoretical approach that uncovers a previously undocumented genome-wide signature of selection in the collective divergence of allele frequencies over time. Applying our approach to temporally resolved allele frequency measurements from laboratory and wild *Drosophila* populations, we quantify the selective contribution to allele frequency divergence and find that selection has substantial effects on much of the genome. We further quantify the magnitude of the total selection coefficient (a measure of the combined effects of direct and linked selection) at a typical polymorphic locus, and find this to be large (of order 1%) even though most mutations are not directly under selection. We find that selective allele frequency divergence is substantial at intermediate allele frequencies, which we argue is most parsimoniously explained by positive — not purifying — selection. Thus, in these populations most mutations are far from evolving neutrally in the short term (tens of generations), including mutations with neutral fitness effects, and the result cannot be explained simply as a purging of deleterious mutations.

**Author summary:** Natural selection is the process fundamentally driving evolutionary adaptation; yet the specifics of how natural selection molds the genome are contentious. A prevailing neutralist view holds that the evolution of most mutations is essentially random. Here, we develop new theory that looks past the stochasticity of individual mutations and instead analyzes the behavior of mutations across the genome as a collective. We find that selection has a strong non-random influence on most of the *Drosophila* genome over short timescales (tens of generations), including the bulk of mutations that are not themselves directly targeted by selection. We show that this likely involves ongoing positive selection.

## Introduction

One of the central problems of evolutionary biology is to delineate the role of natural selection in shaping genetic variation. Most genetic variation consists of neutral mutations which, though having no appreciable effects on fitness, are not free from the influence of selection. When selection acts on non-neutral mutations, neutral mutations that share similar genetic backgrounds can be dragged along for the ride, a process called linked selection [1]. The extent to which linked selection influences neutral variation is a major point of contention [2, 3] — one with practical implications because putatively neutral mutations are widely used to infer population demographic history [4] and as a baseline for detecting selection [2, 5]. There is also ongoing debate about the particular modes of selection responsible for shaping genetic variation. Negative selection purging the influx of deleterious mutations is probably prevalent [6, 7], but positive selection on rarer advantageous mutations is crucial for adaptive evolution and likely also has a hand in shaping neutral variation [8].

Until recently, the bulk of the evidence entering the above debates rested on patterns of genetic variation measured at single snapshots in time. The interpretation of such evidence is complicated because the prospective signatures of selection are accumulated over an uncertain history during which other confounding processes (e.g. population demography) also shape genetic diversity [5, 9, 10]. Crucially, single snapshot data is unable to reveal what the process of selection is doing at any point in time i.e. selectively changing allele frequencies.

A more direct approach is to analyze allele frequency data gathered from the same population at multiple points in time [10]. Evolve and resequence (E&R) experiments [11–14] and studies on wild populations [15, 16] have identified allele frequency changes associated with rapid phenotypic adaptation. However, determining the full nature and extent of selective allele frequency change has been difficult. Numerous methods exist for inferring selection coefficients from allele frequency time series [10, 17–24], but are only reliable for selection that is strong relative to the intensity of random, non-selective allele frequency change (random genetic drift). This is a major limitation that likely precludes detection of most of the influence of selection. Fitness-relevant traits are often complex (influenced by a large number of genes) and harbor ample genetic variation. Selection on such traits will thus often cause modest allele frequency shifts distributed across many loci rather than be concentrated at a small number of strongly selected loci [25–27]. Moreover, even if some genomic regions harbor strongly selected alleles, much of the associated linked selection could be undetectably weak. Thus, resolving the short-term (∼ tens of generations) influence of selection across the genome remains an important challenge [28].

Here we present a new approach to analyze the genome-wide influence of selection using time-resolved allele frequency data. Our approach capitalizes on a distinctive pattern of among-locus temporal allele frequency divergence that to our knowledge has not previously been described. In contrast with single-locus approaches, this allele frequency divergence is a collective pattern incorporating alleles across the genome. We therefore lose the ability to identify particular loci under selection; in return are able to detect polygenic selective processes that are not detectable with single-locus approaches.

Traditionally the allele frequency variance in a cohort of neutral alleles with initial frequency *p* is assumed to have the binomial form

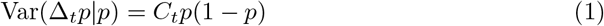

where Δ_*t*_*p* denotes the change in allele frequency after *t* generations, and the variance coefficient *C*_*t*_ is frequency independent [29, Chap. 3]. The allele frequency divergence in Eq. (1) is largely a consequence of random genetic drift. However, selection can also cause neutral allele frequencies to diverge. The influential effective population size literature has derived (frequency-independent) expressions for *C*_*t*_ in a wide variety of circumstances [30]. Crucially, a large body of work has attempted to subsume the effects of selection on neutral alleles into the frequency-independent value of *C*_*t*_, including both the effects of unlinked fitness variation [31, 32], and some manifestations of linked selection [6, 33, 34]. The effective population size literature thus views (1) as a broadly applicable model of neutral allele divergence, simply requiring a tuning of *C*_*t*_ to capture the effects of selection on neutral alleles, at least to a first approximation [3, 30].

Here we show, on the contrary, that linked selection causes among-locus neutral allele frequency variance to deviate from the binomial form (1), such that the variance coefficient *C*_*t*_ is frequency-dependent. We use this frequency-dependence to detect the presence of selection, analyze its influence on allele frequencies over time and estimate the typical magnitude of total selection coefficients (capturing both direct and linked selection) across the genome. Applying our approach to E&R and wild *Drosophila* single nucleotide polymorphism (SNP) data we find evidence of strong linked selection affecting most SNPs (although we cannot rule out migratory fluxes in the wild population). We argue that the specific form of frequency-dependence we find implies a substantial role for positive selection.

## Results

### Neutral evolution implies binomial allele frequency variance

The Eq. (1) binomial variance classically arises in the neutral Wright-Fisher model, which assumes random sampling of gametes each generation; then *C*_*t*_ = 1 − (1 − 1*/*2*N*)^*t*^ where *N* is the (diploid) population size. In its basic form the neutral Wright-Fisher model entails a number of biological simplifications including random mating, constant *N*, non-overlapping generations, and the absence of fitness differences between individuals. Many of these assumptions can be relaxed without affecting the binomial form of Eq. (1), at least approximately for large *N* and over long timescales [30, 35]. Similarly, much of the justification for Wright-Fisher as a biologically valid description of genetic drift is derived from its equivalence to a broader class of drift models in the limit of large *N* and slow allele frequency change (the diffusion limit [36]). Here we are interested in shorter time scales (≤ tens of generations), and want our approach to be applicable to small laboratory populations (*<* 10^3^ individuals). We therefore evaluate the validity of Eq. (1) more generally.

An enormous variety of purely neutral genetic drift models have binomial variance [37]. This includes the Cannings model, which represents neutrality using a general exchangeability assumption that allows for arbitrary offspring number distributions [38]. Binomial variance thus accommodates fundamental deviations from Wright-Fisher such as “sweepstakes” reproduction in high-fecundity organisms [39]. However, due to the presence of fitness variation in adapting populations, neutral mutations do not evolve according to “pure drift” of the sort studied in ref. [37], even if unlinked from alleles under selection [31]. In particular, the Cannings model is not applicable because exchangeability precludes fitness variation between individuals.

In Supplement S1, we show that binomial variance applies quite generally for neutral alleles unlinked from selected loci. In short, we use a generalized exchangeability argument to show that binomial variance holds in the presence of fitness variation provided that the neutral alleles under consideration are in linkage equilibrium with alleles under selection. Intuitively, linkage equilibrium ensures that the distribution of genetic backgrounds is exchangeable between alternate neutral alleles, even though individual genetic backgrounds are not exchangeable.

Non-binomial variance (equivalently, frequency-dependent *C*_*t*_) thus signifies a violation of generalized exchangeability. The obvious way for this to occur is for allele frequency change to have a nonzero bias; this could be due to linked selection, migration, mutation bias or gene drive. Additionally, deviations from binomial variance can occur if the population is structured into genetically differentiated demes (Supplement S2). Below we check for binomial variance empirically and discuss our findings in relation to these factors leading to non-binomial variance, focusing mostly on selection for reasons that will become apparent.

Note that while our exchangeability argument yields Eq. (1) with finite variance for finite *N*, in the diffusion limit infinite variance is possible in the Cannings model [37]. None of our results depend on *N* → ∞ limiting behavior, so we do not discuss this possibility further.

### Selection creates non-binomial allele frequency variance

We now analyze the effects of selection on allele frequency divergence, demonstrating that deviations from binomial variance will often result.

The expected frequency change after one generation due to selection on an allele starting at frequency *p* is given by

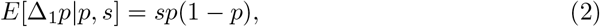

where 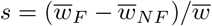 is the selection coefficient, 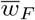 is the mean fitness of the focal allele, 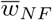 is the mean fitness of all other alleles at the same locus, and 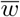 is population mean fitness. Here *s* is the “total” selection coefficient that captures the net effect of selection at linked loci and the focal locus [40, 41].

Selection generates among-locus divergence of allele frequencies when its strength or sign varies among alleles in a cohort. To quantify this effect, we apply the law of total variance to Δ_1_*p* where *s* is allowed to vary between loci:

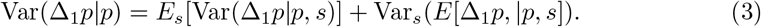

The second term in Eq. (3) represents the deterministic allele frequency divergence created by among-locus variation in *s*. Using Eq. (2), it can be written as *σ*^2^(*s*|*p*)[*p*(1 − *p*)]^2^, where *σ*^2^(*s*|*p*) is the (possibly frequency-dependent) variance in total selection coefficients among loci with initial frequency *p*. The presence of the [*p*(1 − *p*)]^2^ factor will tend to cause intermediate frequency alleles to have elevated variance relative to the binomial case. Thus, while it is possible for the allele frequency variance created by selection to be binomial, in general it is not. Beyond the tendency for elevated variance at intermediate frequencies, the exact shape and magnitude of the deviation is determined by *σ*^2^(*s*|*p*).

More generally, allele frequencies are observed *t >* 1 generations apart during which time the selective divergence accumulates. Assuming that the selective allele frequency change at most loci is small over the measurement interval (at most loci in the cohort, 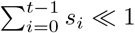 and |*s*_*i*_| ≪ 1, where *s*_0_, …, *s*_*t−*1_ are total selection coefficients in the intervening generations), the selective divergence generalizes to 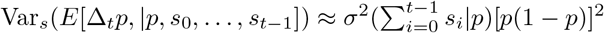, which now depends on the temporal structure of selection. Sustained selection, which manifests as positive among-locus covariances between the *s*_*i*_, causes rapid divergence with allele frequency variance growing quadratically over time if selection coefficients are constant (Supplement S3). Randomly fluctuating selection also creates divergence, but with variance accumulating more slowly at a linear rate — a selective random walk [42]. Selection that fluctuates in a more predictable manner could in principle generate no divergence at all e.g. if the loci in a cohort share a cyclical pattern.

The selective divergence just described is central to our results. Selection has another effect in Eq. (3) that is not important to our results: it perturbs the drift contribution to divergence *E*_*s*_[Var(Δ_*t*_*p*|*p, s*_0_, …, *s*_*t−*1_)]. This effect occurs when a mean selective bias in the cohort displaces allele frequencies and thus perturbs the effects of drift (regardless of whether there is among-locus variation in total selection coefficients). In Supplement S4 we show that the selective perturbation to the drift variance has the form 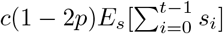 where *c* is a frequency-independent constant of order 1. This result assumes that the cohort does not start close to fixation, and is also insensitive to population dynamic specifics if many generations separate measurements (*t* ∼ 10 in the data we analyze). For a generational measurement interval (*t* = 1) this result holds in canonical models (i.e. Wright-Fisher and Moran), but in general it is possible that the exact form of the selective drift perturbation depends on population specifics. In the following analysis the exact expression for the selective drift perturbation will not be important; we only use the fact that it scales with 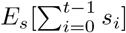, which implies that its effects are negligibly small in the populations of interest here (Methods).

Combining variance contributions we have

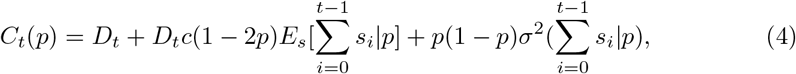

where *D*_*t*_ is the frequency-independent variance coefficient in the absence of selection. The variance coefficient *C*_*t*_(*p*) is thus partitioned respectively into a frequency-independent genetic drift component, a frequency-dependent selective drift perturbation, and a frequency-dependent selective divergence.

### Intermediate frequency alleles have elevated variance in *Drosophila*

In light of the preceding results, we investigated whether binomial allele frequency variance is observed empirically. To check for binomial frequency dependence, we calculated *C*_*t*_ = Var(Δ_*t*_*p*|*p*)*/p*(1 − *p*) using SNP allele frequencies measured in two sequencing snapshots, binning SNPs according to their initial frequency *p*.

In two fruit fly (*D. Simulans*) E&R experiments [11, 12], we observe systematically elevated *C*_*t*_ at intermediate frequencies (Fig. 1). We rule out measurement error as driving this pattern, because the major sources of pooled sequencing error (population sampling, read sampling, unequal individual contributions to pooled DNA) also create binomial variance rather than a systematic frequency-dependent bias (Supplement S5; [43, 44]). We also rule out migration, since these E&R populations are closed. Moreover, as will be discussed in the next section, systematically elevated variance cannot be explained by a few large effect loci, implying that a substantial fraction of SNPs across the genome are involved in the observed pattern. Hence we also rule out mutation bias and gene drive as being the main driver of elevated variance at intermediate frequencies since these processes do not have the requisite scale. Finally, population structure tends to create a variance deficit at intermediate frequencies (Supplement S2); thus, even if some population structure is present in these closed E&R populations, it would tend to eliminate the observed elevation of variance, not explain it. We deduce that the pattern observed in Fig. 1 is due to selection, consistent with the theoretical prediction that the selective divergence tends to cause elevated variance at intermediate frequencies.

**Fig 1.**
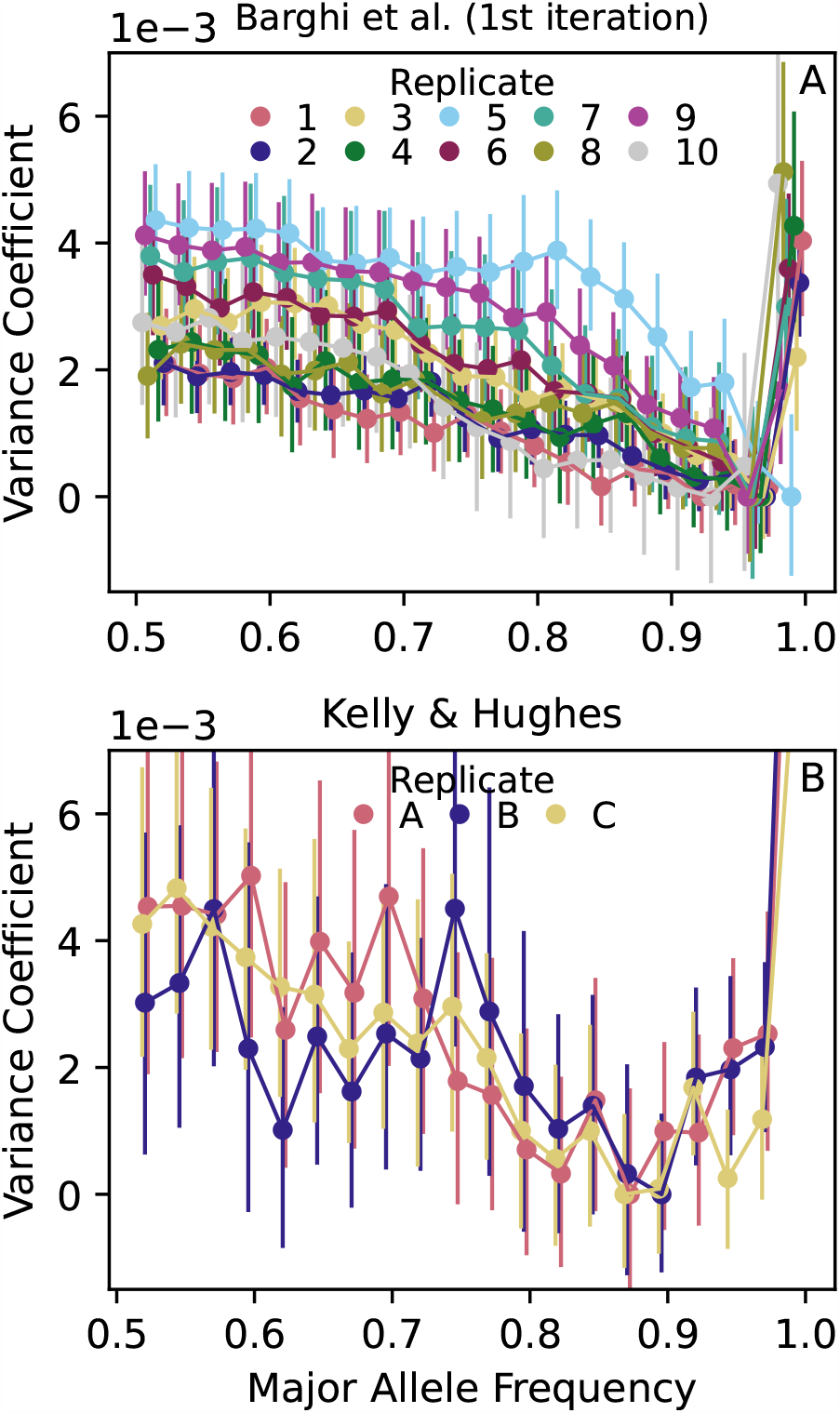
Intermediate frequency SNPs in E&R *D. Simulans* populations (A [11]; B [12]) have systematically elevated variance coefficients *C*_*t*_(*p*) = Var(Δ_*t*_*p*|*p*)*/p*(1 − *p*) relative to higher frequency SNPs after one round of evolution and resequencing (*t* ≈ 10 in A; *t* ≈ 15 in B), inconsistent with the binomial expectation for neutrally evolving alleles (1). *C*_*t*_(*p*) is calculated in 2.5% major allele frequency bins using all SNPs in the genome (circles). Vertical lines show 95% block bootstrap confidence intervals (1Mb blocks). We subtract the constant min_*p*_*C*_*t*_(*p*) from *C*_*t*_(*p*) in each replicate to prevent differences in the overall magnitude of *C*_*t*_(*p*) between replicates from obscuring *p* dependence within each replicate.

Similar results are found in a wild *D. Melanogaster* population [15] (not shown in Fig. 1; see Fig. 2B), although this population is not closed and elevated variance could also be attributed to migration. The effect of migration on allele frequency divergence can be understood analogously to selection (Eq. (3)) as introducing a migration divergence term Var(*m*(*p*^*∗*^ − *p*)|*p*) = *m*^2^Var(*p*^*∗*^ − *p*|*p*) where *m* is the proportion of individuals in the focal population replaced by migrants from the source population each generation, and *p*^*∗*^ denotes source population frequencies. The migration divergence thus depends on the structure of differentiation between focal and source populations. The *a priori* expectation is for Var(*p*^*∗*^ − *p*|*p*) to be greatest at high *p* (the opposite of the observed pattern), where the largest differences *p*^*∗*^ − *p* are possible (analogous to the mathematical constraints on *F*_*ST*_ [45]). However, since we do not know the structure of population differentiation (or even what the source population might be), we remain agnostic about the influence of migration in the ref. [15] population.

**Fig 2.**
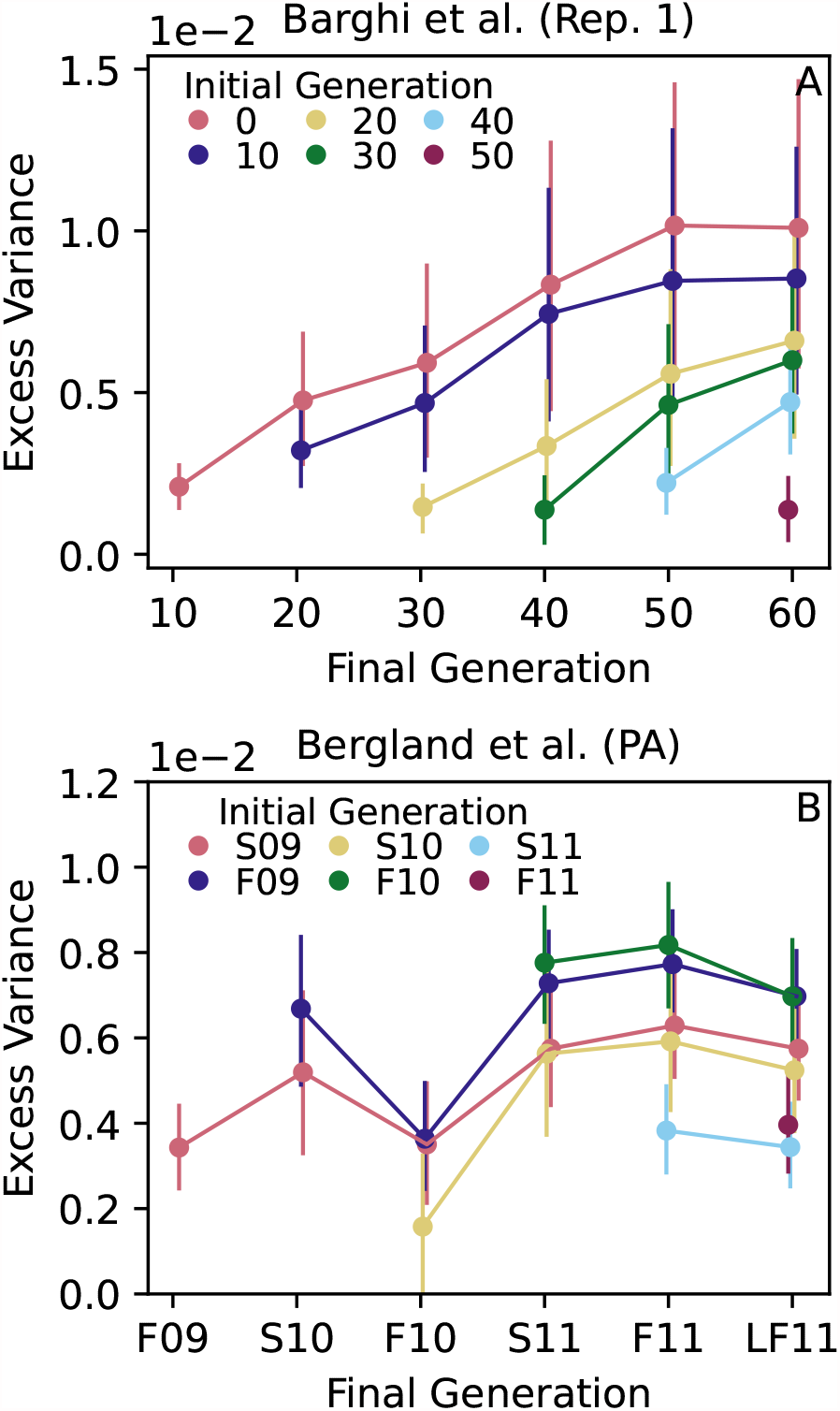
Excess allele frequency variance (a measure of deviation from neutrality defined as *C*_*t*_(*p*) −*C*_*t*_(*p*^*∗*^)) accumulates over time in a *D. Simulans* E&R experiment (A; [11]), but remains relatively flat in a wild *D. Melanogaster* population (B; S=Spring, F=Fall, LF=Late Fall; 09=2009 etc.; [15]). The excess variance is calculated for alleles falling within a 2.5% major allele frequency bin at *p* = 0.5 (i.e. 0.475 *< p <* 0.525), and similarly for the reference variance *C*_*t*_(*p*^*∗*^) with *p*^*∗*^ = 0.9 (A) and *p*^*∗*^ = 0.8 (B). Vertical lines show 95% block bootstrap confidence intervals (1Mb blocks).

Next we explored the behavior over time of the elevated variance shown in Fig. 1 by following its accumulation within a frequency cohort for the two studies in which allele frequencies were measured more than twice [11, 15]. At each measured timepoint we quantified the excess variance using the difference *C*_*t*_(*p*) − *C*_*t*_(*p*^*∗*^), where *p* is the initial frequency of the cohort and *p*^*∗*^ *> p* is a reference frequency. In practice we choose *p* = 0.5 to maximize the contrast with the reference frequency, while *p*^*∗*^ ∼ 0.8 − 0.9 is chosen to be large enough that there is a meaningful contrast with *p* = 0.5 but safely displaced from the *p* = 1 boundary where allele frequency variances are not measured reliably (see sharp increases in Fig. 1 as *p* → 1).

We find that excess variance accumulates over the course of the entire Barghi et al. [11] E&R experiment (Fig. 2A shows one replicate, other replicates are similar; Fig. S1), implying a sustained, polygenic divergence in allele frequencies. This pattern is consistent with the positive Δ*p* temporal autocovariances documented in [28]. Sustained divergence is what we expect to occur from selection in a novel but constant laboratory environment.

By contrast, excess variance in wild *D. Melanogaster* populations [15] does not exhibit continual accumulation of excess variance over time, with fluctuations evident in each cohort (Fig. 2B). Fluctuations imply a concurrent reversal in the direction of non-neutral allele frequency change across many loci such that non-neutral divergence is partly lost to a subsequent coordinated non-neutral convergence. Bearing in mind that migration may contribute to this pattern, the fluctuations shown in Fig. 2 are compatible with temporally fluctuating selection affecting a large proportion of the genome, as proposed by ref. [15]. However, while ref. [15] attributed temporal fluctuations in selection to periodic seasonal change, we do not see a clear annual periodicity in the accumulation of variance. A similar lack of annual periodicity is found in the corresponding temporal autocovariances [28]. These results suggest a more complex selective (or migratory) regime of which seasonal fluctuations are only a part.

### Linked selection strongly perturbs SNP frequencies in *Drosophila*

In the previous section we argued that selection is most likely responsible for elevated allele frequency divergence at intermediate frequencies in three *Drosophila* studies (with the possible exception of the ref. [15] study because of migration). We next used the theory developed above to estimate the typical magnitude of total selection coefficients associated with elevated divergence (we also apply our analysis to ref. [15] supposing that selection was responsible).

We measure the typical intensity of selection using the among-locus standard deviation *σ*(*s*|*p*). This quantity determines the selective divergence in Eq. (4), and has the convenient property of measuring the absolute magnitude of *s* regardless of sign. Intuitively, *σ*(*s*|*p*) measures the intensity of a collective “polygenic” adaptive response shared across many loci. If a fraction *f* of loci have *s* = 0, then 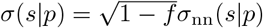 where *σ*_nn_(*s*|*p*) is the standard deviation in *s* among non-neutral loci. Thus, a substantial fraction of the alleles in a cohort must have nonzero *s* (*f* appreciably smaller than 1) for there to be a discernible *σ*(*s*|*p*) signal.

We estimate *σ*(*s*|*p*) from measured allele frequency divergence using Eq. (4). Since we only have measurements separated by *t* generations, we actually estimate 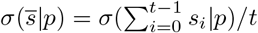 where 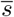 is the time-averaged selection coefficient 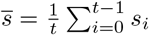. To estimate 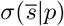 from Eq. (4), we need to eliminate the non-selective divergence contributions of genetic drift *D*_*t*_ and measurement error (which was not included in Eq. (4)). In Methods we show that these latter contributions are cancelled out in the excess variance *C*_*t*_(*p*) − *C*_*t*_(*p*^*∗*^), avoiding the complication of independently estimating them. However, some selective divergence is also cancelled out in the difference *C*_*t*_(*p*) − *C*_*t*_(*p*^*∗*^), so that this approach only obtains a lower bound

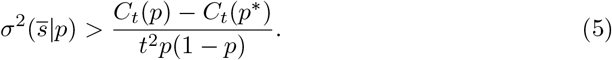

In all three *Drosophila* studies, we find the above lower bound to be of order 10^*−*4^ (Fig. 3), implying that total selection coefficients with magnitudes of order 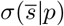 ∼ 1% are commonplace in the populations considered here.

**Fig 3.**
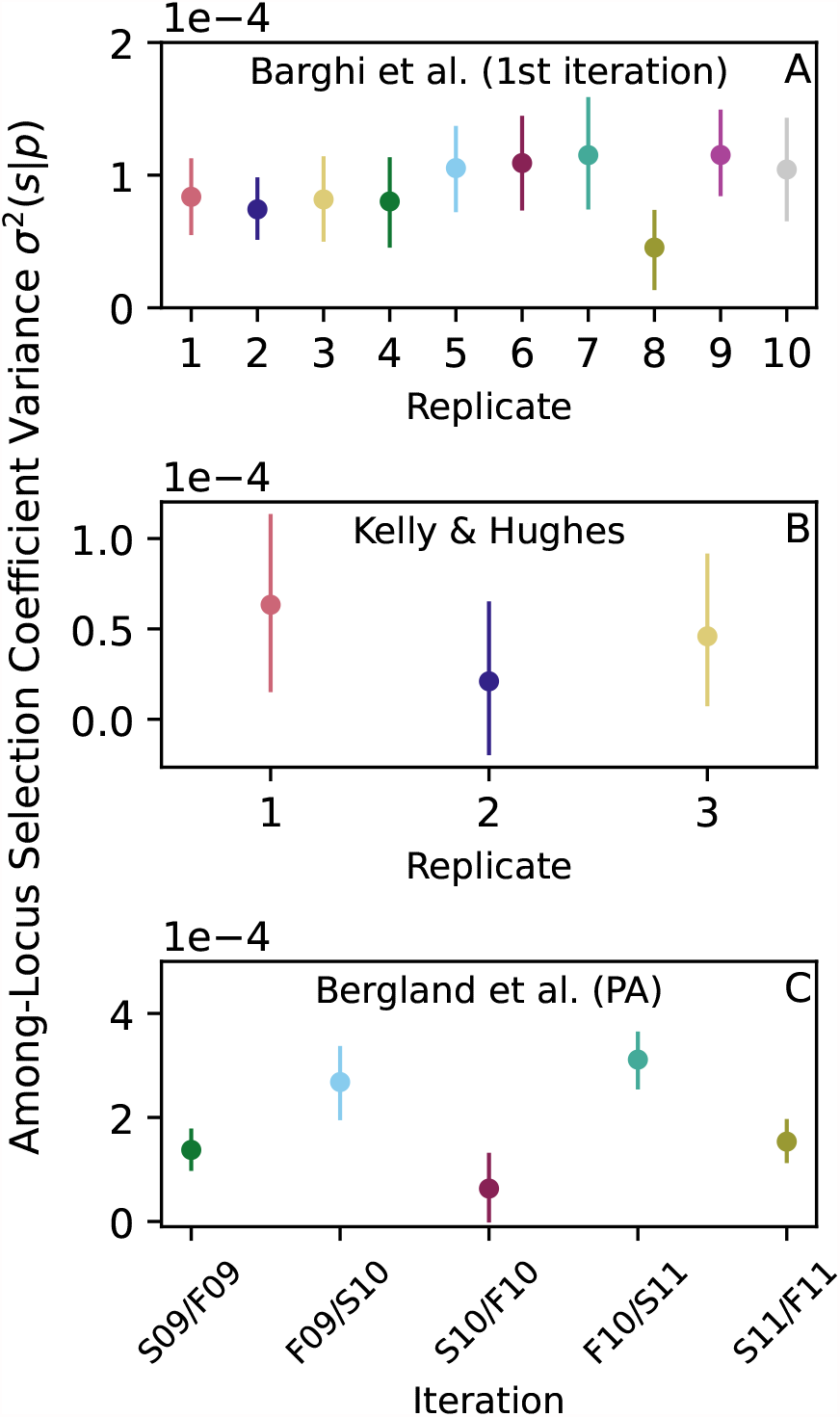
Total selection coefficients show substantial among-locus variance in *Drosophila*. (A-C) Lower bound estimates of 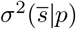 calculated from (5) (circles; vertical lines show 95% block bootstrap confidence intervals) are of order 10^*−*4^, which implies typical *s* values of ∼1%. Following the original studies [11, 12, 15], we assume *t* = 10 (A); *t* = 15 (B) and *t* = 10 (C; for both summer and winter).

### Positive selection explains consistently positive excess variance

In the preceding sections we showed that selection strongly influences temporal allele frequency divergence throughout the *Drosophila* genome. In principle this could involve a mix of positive and negative selection, although *a priori* we might expect that the presence of such pervasive selection is more easily explained by negative selection acting continually to purge deleterious mutations. However, it is not clear how a negative selection scenario can be reconciled with the observed pattern of elevated variance coefficients at intermediate frequencies (Fig. 1). The effects of negative selection should be substantially stronger at high major allele frequencies, because deleterious mutations strong enough to cause detectable allele frequency divergence are unlikely to reach high frequencies.

To better understand the processes generating excess variance we performed forward-time population genetic simulations of negative and positive selection scenarios. We find that negative selection indeed does not consistently generate positive excess variance (Fig. 4A), because 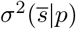 rapidly increases with *p* such that the overall selective divergence term 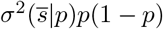 in Eq. (4) is independent of frequency — just like a neutral allele frequency divergence. The effects of positive selection on the other hand are elevated at intermediate frequencies leading to consistently positive excess variance (Fig. 4A,C). Some amount of positive selection therefore seems to be essential to explain the elevated divergence in Figs. 1-3.

**Fig 4.**
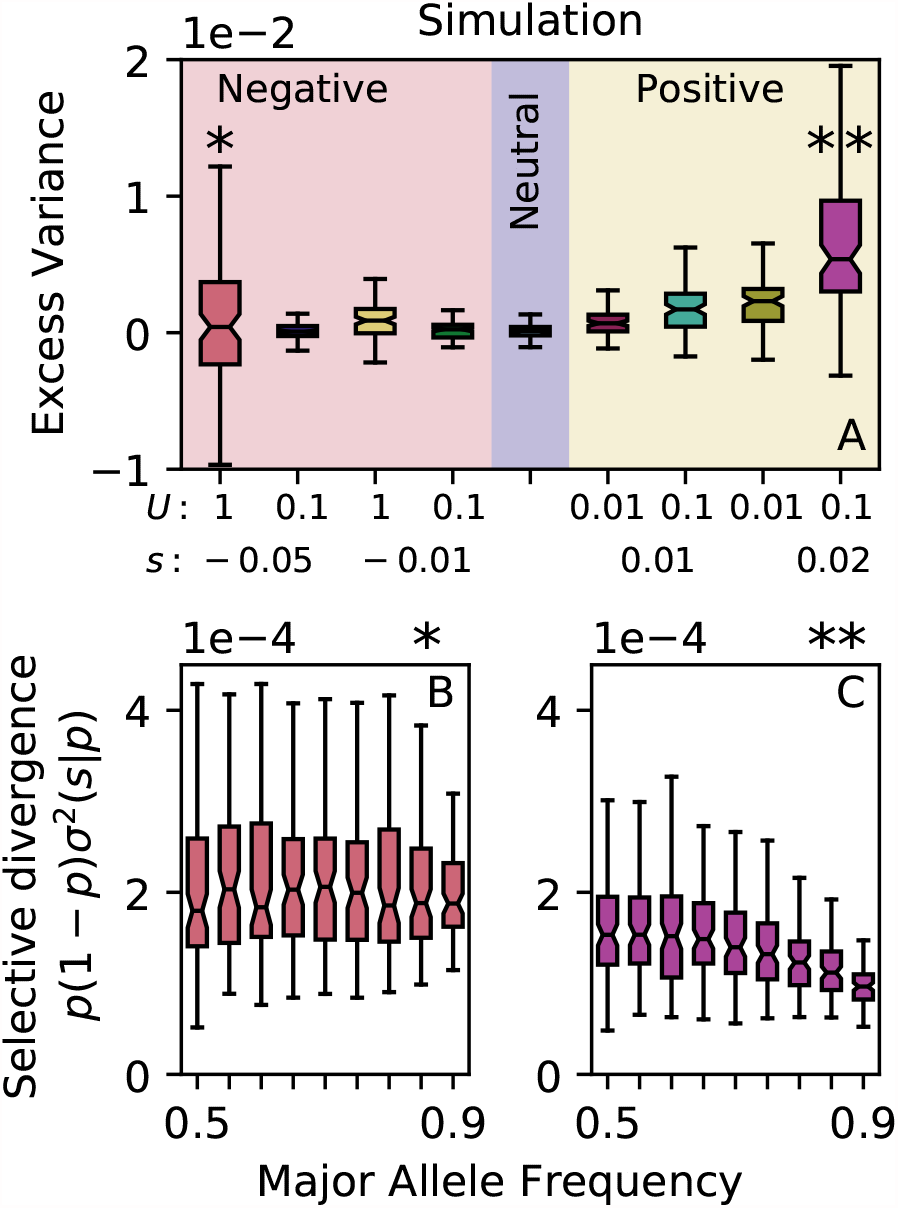
(A) Forward-time population genetic simulations only consistently show elevated excess variance under positive selection. Excess variance defined as *C*_*t*_(*p*) − *C*_*t*_(*p*^*∗*^) with major allele frequencies 0.5 *< p <* 0.55 and 0.9 *< p*^*∗*^ *<* 0.95 and *t* = 10 generations. (B) Under strong negative selection (deleterious mutation rate *U* = 1/genome/generation, mutation selection coefficient *s* = −0.05), total selection coefficients are substantial at all frequencies but much stronger for high major allele frequencies resulting in a frequency-independent overall selective divergence 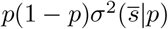 like the neutral case. (C) In contrast, the selective divergence 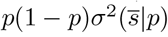 shows clear frequency dependence under positive selection, thus producing excess variance at intermediate frequencies. Population size *N* = 1000; 100 replicates per parameter combination.

## Discussion

Several lines of evidence support the view that selection strongly influences genetic variation in *Drosophila* [8, 12, 28, 46]. Our results independently show that even over a short time interval (tens of generations), most intermediate frequency SNPs are influenced by selection — total selection coefficients (which include linked selection) of |*s*| ∼ 1% are the norm among intermediate frequency SNPs, despite most of these SNPs having no effect on fitness. Since our method relies on contrasting behavior at different frequencies, the effect of selection on extreme frequency alleles is used as a reference and is therefore not directly inferred. We expect the effects of selection to be even greater at extreme frequencies where most deleterious mutations are segregating and recent neutral mutations are most tightly linked to selected backgrounds.

The power of our approach stems from aggregating allele frequency behavior over many loci, thereby leveraging the sheer number of variants measured with whole-genome sequencing to discern a selective signal. Heuristically, the sampling error in the lower bound estimate (5) is proportional to 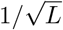 where *L* is the number of independent loci used to estimate *C*_*t*_(*p*). With enough sequenced variants (*L* ∼ 10^5^), selection coefficients of order |*s*| ∼ 1% should be detectable over a single generation even when allele frequency noise is of comparable magnitude (i.e. read depth and population size ∼ 10^2^; see Methods). Intuitively, variants across the genome experience a detectable non-neutral shift as a collective even though the underlying allele frequency changes may be indistinguishable from drift at individual loci.

Our approach is a departure from the widespread use of frequency-independent *C*_*t*_ for neutral mutations [30]. The variance coefficient *C*_*t*_ can be expressed in terms of the “variance effective population size” *N*_*e*_ as *C*_*t*_ = 1 − (1 − 1*/*2*N*_*e*_)^*t*^. Thus, selection makes *N*_*e*_ frequency-dependent for neutral mutations over short timescales (i.e. before an appreciable fraction of the alleles in a cohort fix). The origin of this non-binomial allele frequency variance is variation in the selective background of alleles at different loci.

Selection does not need to be consistent over time to have this effect: stochastically fluctuating selection with no temporal consistency will also generate non-binomial allele frequency variance. However, temporally consistent selection generates divergence more rapidly, and temporal covariances can be responsible for most of the selective divergence (Results; Supplement S3). Moreover, allele frequency changes Δ*p* are correlated over time in the systems analyzed here [28]. Thus, it seems likely that temporally consistent selection is at least partly responsible for the patterns documented here.

Note, however, that in contrast to ref. [28], the temporal covariances relevant to allele frequency divergence are between total selection coefficients, not Δ*p* (Supplement S3). Covariances in total selection coefficients are only non-zero if those coefficients vary among loci, whereas Δ*p* covariances quantify any temporal consistency in allele frequency change [47]. Thus, Δ*p* temporal covariances can theoretically be present without any selective divergence, and vice versa. In practice, the temporal autocovariances in Δ*p* must be calculated across three measurement steps e.g. Cov(*p*_*t*_ − *p*_0_, *p*_2*t*_ − *p*_*t*_). These cross-measurement covariances do not contribute to the divergence observed at *t* generations, and are only a subset of the covariances contributing to the divergence observed at 2*t* generations. Therefore, the patterns of variance accumulation documented here are related but not equivalent to the patterns documented in ref. [28]. Temporal autocovariances in Δ*p* predominantly capture the extent to which the genome-wide influence of selection has a temporally enduring pattern across measurements. Allele frequency divergence captures the cumulative genome-wide influence of both temporally stable and fluctuating selection between two measurements. The relative contribution from temporal covariances in total selection coefficients depends on the intensity of selective fluctuations as well as the persistence time of linkage disequilibrium (Supplement S3), and would require generational allele frequency measurements to quantify.

We found that the frequency structure of allele frequency divergence is informative about the underlying structure of direct selection (Fig. 4). Elevated divergence of intermediate frequency alleles is difficult to explain if only negative selection on deleterious mutations is occurring. Although selection against deleterious mutations can generate transient sweep-like behavior for neutral mutations that originate on genetic backgrounds with above-average fitness, negative selection has overwhelmingly more influence on allele frequency dynamics at low/high frequencies compared to intermediate frequencies (Fig. 4; [48]). More broadly, it may be possible to make more detailed inferences about the structure of direct selection by moving beyond allele frequency variances and analyzing the entire distribution of allele frequency change Δ_*t*_*p*.

Quantifying the bounds on how much selection is possible, and how much selection actually occurs in natural popoulations, is a long running controversy [49, 50]. The strong total selection coefficients (|*s*| ∼ 1%) we find must predominantly reflect linked selection on neutral SNPs. This implies a substantial risk of overestimating the amount of direct selection when, as is commonly done, selection coefficients are inferred at individual loci and then attributed to direct selection. This “excess significance” is a well known difficulty in E&R experiments [12, 51], and similar challenges have arisen in wild populations [15]. Our results indicate that improving the sensitivity of single-locus selection coefficient inferences, or better controlling for multiple comparisons, will likely not resolve this issue. Our total selection coefficient estimates are also substantially larger than direct selection coefficients of individual alleles estimated from diversity patterns in *Drosophila* [8]. This is consistent with a linkage-centered view of neutral mutation evolution in which the selective background of most neutral mutations contains multiple alleles under selection such that allele frequency behavior is governed by the fitness variation within local “linkage blocks” [52] or larger haplotypes [11].

## Methods

### Data sources and availability

All processing and plotting code can be accessed at https://github.com/jasonbertram/polygenic_variance/. SNP frequency data were obtained from the open access resources published in [15] (wild *D. Melanogaster*, 1 replicate, ∼ 5 × 10^5^ SNPs, 7 timepoints), [11] (*D. Simulans* E&R, 10 replicates, ∼ 5 × 10^6^ SNPs, 7 timepoints) and [12] (*D. Simulans* E&R, 3 replicates, ∼ 3 × 10^5^ SNPs, 2 timepoints). We performed no additional SNP filtering. For the [15] data, only SNPs tagged as “used” were included.

### Block bootstrap confidence intervals

We use bootstrapping to estimate the variability of the quantities plotted in Figs. 1-3. These quantities are calculated as an average over loci, where nearby loci are unlikely to be statistically independent due to linkage. To account for the non-independence of individual loci when bootstrap sampling, 95% confidence intervals are calculated using a block bootstrap procedure [28]. Each chromosome is partitioned into 1 megabase windows (∼ 120 total windows). Bootstrap sampling is then applied to these windows. The plotted vertical lines span the 2.5% and 97.5% block bootstrap percentiles.

### Estimation of the selection coefficient variance

To derive Eq. (5) we show that the reference value *C*_*t*_(*p*^*∗*^) satisfies the inequality

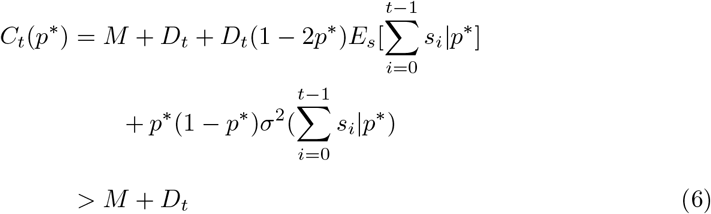

The first line above is Eq. (4) evaluated at the reference frequency *p*^*∗*^ with an additional measurement error term *M* included. *M* is frequency-independent because measurement error is binomial (Supplement S2; [43, 44]). Eq. (6) implies that the reference value *C*_*t*_(*p*^*∗*^) is an upper bound on the drift and measurement components of *C*_*t*_(*p*) for all *p*. Taking the difference *C*_*t*_(*p*) − *C*(*p*^*∗*^), we then have

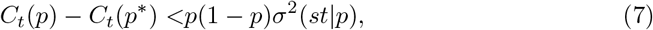

eliminating *D*_*t*_ and *M*.

To derive Eq. (6) we first eliminate the selective drift perturbation 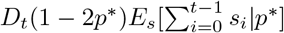. This term is negligibly small compared to the selective divergence in the populations considered here: *C*_*t*_ (and therefore *D*_*t*_) is of order 10^*−*2^, *E*[*s*_*i*_|*p*] is at most of order 10^*−*2^, and *t* ∼ 10; hence 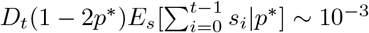. By comparison, 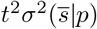 is of order 10^*−*2^. Second, we have *p*^*∗*^(1 − *p*^*∗*^)*σ*^2^(*st*|*p*^*∗*^) *>* 0; subtracting this term gives the inequality.

### Simulations

We used SLiM [53] to simulate a closed population with *N* = 10^3^ individuals, a 100Mb diploid genome, a recombination rate of 10^*−*8^/base pair/generation, and a neutral mutation rate of 10^*−*8^/base pair/generation. Non-neutral mutations were introduced at rate *U* /chromosome/generation, where in each simulation non-neutral mutations were assumed to have the same fixed selection coefficient. Four background selection regimes (*U* = 1, 0.1 × *s* = −0.05, −0.01), one neutral regime (*U* = 0), and four positive selection regimes (*U* = 0.1, 0.01 × *s* = 0.01, 0.02) were evaluated (Fig. 4). In each regime, 100 replicates were simulated with complete genotypes recorded at generations 10^4^ and 10^4^ + 10, mimicking the *t* = 10 generation interval in the empirical studies after a burn in period of 10*N* = 10^4^ generations. Total selection coefficients in Fig. 4 computed using (2) from genotype data at generation 10^4^.

### Estimation limits

Our analysis relies on detecting differences in *C*_*t*_(*p*) between cohorts with different values of *p*. The ability to detect such differences is determined by the sampling error in *C*_*t*_(*p*) arising due to the calculation of Var(Δ_*t*_*p*|*p*) from a finite number of loci. To estimate this sampling error, we assume that Δ_*t*_*p* is approximately normally distributed, in which case the sample variance in Var(Δ_*t*_*p*|*p*) is 2Var(Δ_*t*_*p*|*p*)^2^*/*(*L* − 1) ≈ 2Var(Δ_*t*_*p*|*p*)^2^*/L* where *L* ≫ 1 is the number of independent loci used to estimate Var(Δ_*t*_*p*|*p*). The standard error in *C*_*t*_(*p*) = Var(Δ_*t*_*p*|*p*)*/p*(1 − *p*) is thus given by 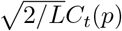. This defines the scale of statistically detectable differences in *C*_*t*_(*p*) − *C*_*t*_(*p*^*∗*^), which in turn determines the statistically detectable lower bound estimate on *σ*^2^(*s*|*p*) (5). For example, to detect *σ*^2^(*s*|*p*) ∼ 10^*−*4^ at *p* = 0.5 (i.e. a typical selection coefficient of *σ*(*s*|*p*) ∼ 1%) after one generation of evolution with *C*_1_ ∼ 10^*−*2^ (i.e. a population sample of ∼ 100 individuals, an average read depth of ∼ 100 and fairly strong genetic drift *D*_1_ ∼ 10^*−*2^), we need at least *L* ∼ 10^5^ independent SNPs.

## Supporting information

**S1 Supplement Supplemental analysis and figures**.

## Acknowledgments

We thank Matthew Hahn for insightful discussions.

## Supplement

### S1 Frequency dependence of *C* for neutrally evolving alleles

Here we show that the allele frequency variance of neutrally evolving alleles has the form Var(Δ_*t*_*p*|*p*) = *C*_*t*_*p*(1 −*p*) where Δ_*t*_*p* is the change in allele frequency over *t* generations, the variance is evaluated across loci with the same initial allele frequency *p*, and the variance coefficient *C*_*t*_ is independent of *p*. Neutrally evolving alleles are defined as having no effect on fitness and being in linkage equilibrium with alleles that do affect fitness.

Our argument is based on a generalization of the Cannings model [1]. Let *y*_*i*_ be the number of copies of an allele descended from the *i*’th copy of that allele *t* generations ago. There are a total of 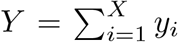 copies of the allele in the population, where *X* is the number of allele copies present *t* generations ago. The allele frequency change that occurs is 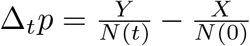, where *N* (0) and *N* (*t*) denote the census population size *t* generations ago and at the present time, respectively. To simplify our equations we assume a haploid population; for a diploid population we replace *N* → 2*N* without otherwise affecting our argument.

Since all loci share the same demography, there is no among-locus variance in *N* (*t*). It follows that the among-locus variance in *Y* for alleles with the same initial frequency *p* (and hence same initial abundance *X*) can be written as 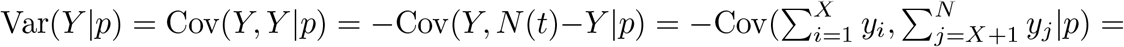 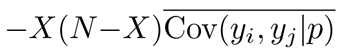, where 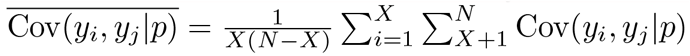 is the descendant number covariance averaged over allele copies. Consequently, we have

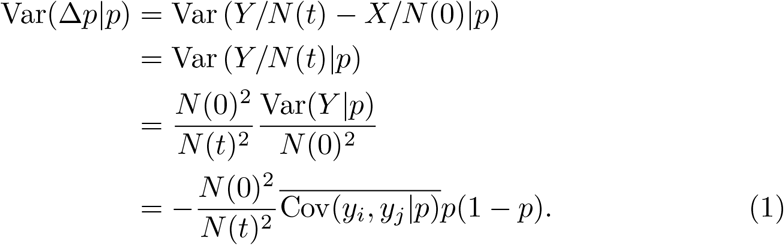

To prove our claim that *C*_*t*_ is independent of *p* we therefore need to show that the averaged covariance 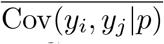 is independent of *p*.

Frequency-independent *C*_*t*_ is straightforward to show in the Cannings model

[2]. Neutrality in the Cannings model takes the form of a symmetry assumption: the random variables *y*_*i*_ are assumed to be exchangeable (i.e. the joint probability distribution of the *y*_*i*_ is invariant with respect to rearrangement of their labels *i*; [3]). Exchangeability allows for a wide variety of marginal distributions for *y*_*i*_ (the allelic offspring distribution), giving the Cannings model enormous biological flexibility (e.g. [2]). When the *y*_*i*_ are exchangeable, 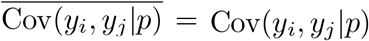 is independent of *i, j* for *i* = 1, …, *X* and *j* = *X* +1, …, *N* because Var(*N* (*t*)) = 0 implies Cov(*y*_*i*_, *y*_*j*_|*p*) = −*σ*^2^*/*(*N* (0) − 1) where *σ*^2^ = Var(*y*_1_) = … = Var(*y*_*N*(0)_) is the variance in the number of allele descendants. This value of Cov(*y*_*i*_, *y*_*j*_|*p*) is independent of *p* because, from the definition of exhcangeability, which allele identities are attached to the *y*_*i*_ has no bearing on their joint probability distribution.

However, the Cannings exchangeability assumption has an important limitation as a model of genetic neutrality, namely that no non-neutral loci can be present in the genome. Non-neutral loci introduce fitness variation among genetic backgrounds which implies differences in the marginal distributions of the *y*_*i*_ corresponding to those genetic backgrounds. Nevertheless, provided that the loci over which Var(Δ_*t*_*p*|*p*) is being calculated are themselves neutral and in linkage equilibrium with loci under selection, 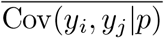 will still be frequency independent. To see this, note first that 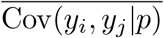 is invariant with respect to reshuffling *i, j* → *i′, j′* of the allele copy labels at each locus:

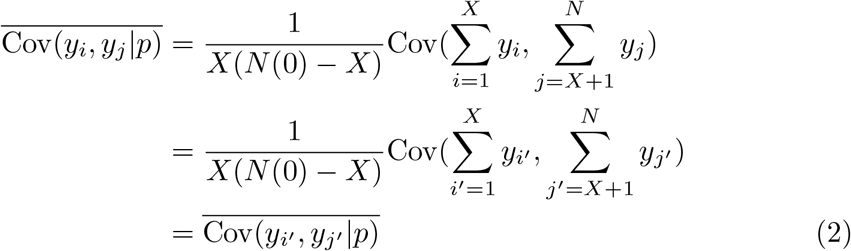

In particular, at each locus we can assign *i, j* labels at random such that *y*_*i*_ and *y*_*j*_ are sampled at random from one focal and one non-focal background respectively. For neutral alleles in linkage equilibrium with non-neutral alleles, the focal and non-focal distributions of genetic backgrounds thus sampled are identical. This leads to a situation analogous to strict exchangeability: the allele identities of the *y*_*i*_ have no bearing on the distribution of pairs *y*_*i*_, *y*_*j*_ that enter the covariance implying that 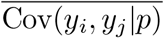 is independent of *p*. This proves our claim.

### S2 Population structure

We assume that the population under consideration consists of *K* subpopulations. Population allele frequencies can be written as a sum over subpopulations 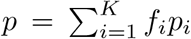,where the *f*_*i*_ are the proportional subpopulation abundances with Σ_*i*_ *f*_*i*_ = 1. Each subpopulation is assumed to evolve neutrally and independently such that Var(Δ_*t*_*p*_*i*_|*p*_*i*_ = *p*_*i*0_) = *C*_*i*_(*t*)*p*_*i*0_(1 − *p*_*i*0_) (Supplement S1) where *p*_*i*0_ is the initial frequency in population *i* and *C*_*i*_(*t*) is independent of *p*_*i*0_. The assumption of independent evolution implies no migration between subpopulations and hence maximal potential for subpopulation differentiation. We discuss the effects of migration further below.

The variance in population Δ_*t*_*p* is evaluated for loci with a fixed initial population *p*, which means that the subpopulation *p*_*i*0_ may differ between loci. Using the law of total variance, we decompose the population variance into terms representing loci with different initial frequency vectors 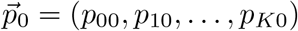,

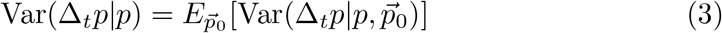

since neutrality within each subpopulation implies 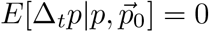. Evaluating the variance inside the expectation gives

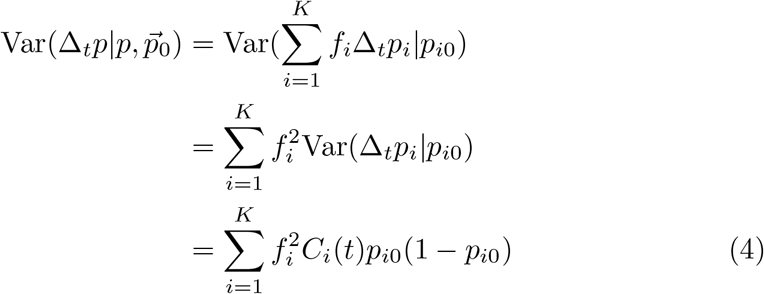

where in the first line we have used the independence of subpopulations (no covariances between the Δ_*t*_*p*_*i*_).

Substituting this expression into (3) and using the fact that 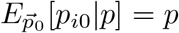, we obtain

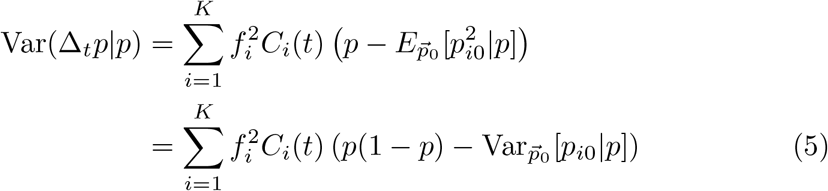

Deviations from binomial variance due to population structure thus arise due to the subpopulation variances 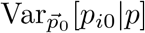, i.e. the among-locus variance in the subpopulation allele frequencies among alleles with population frequency *p* at *t* = 0. In general these variances depend on the structure of genetic variation across subpopulations. To gain some intuition, suppose that the subpopulation-specific variance coefficients *C*_*i*_(*t*) are inversely proportional to census population size such that *f*_*i*_*C*_*i*_(*t*) = *A* has the same value in each subpopulation. We then obtain

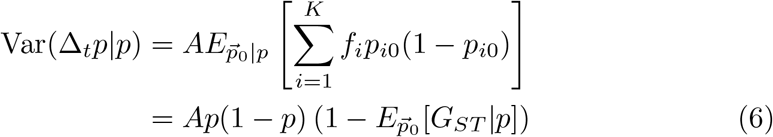

where 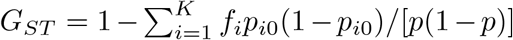 is Nei’s definition of the fixation index at one locus [4]. In the case of a single biallelic locus, *G*_*ST*_ can span almost the entire [0, 1] range at intermediate population frequencies *p*, but approaches zero as *p* → 1 [5]. Since the *G*_*ST*_ term is subtracted in the above equation, this suggests that population structure tends to create a variance deficit at intermediate frequencies compared to a binomial (the opposite of the variance excess described in the main text).

We now drop the *f*_*i*_*C*_*i*_(*t*) = *A* restriction used above to connect our analysis to *G*_*ST*_, and argue that 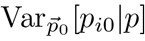 grows faster than binomial as 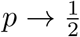 under broadly applicable biological conditions.

We first assume that the allele frequencies in each subpopulation are independent. Since new mutations arise at low frequency and most are rapidly lost before reaching intermediate frequencies, the distribution of allele frequencies (the folded site frequency spectrum) in each subpopulation is strictly decreasing with increasing *p*. The joint distribution of *p*_*i*0_, and the frequency of the same allele in the rest of the population 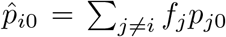, is thus the product of two distributions, *P* (*p*_*i*0_) and 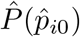, each strictly decreasing with increasing *p*. The variance 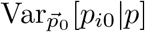 is determined by the distribution of *p*_*i*0_ along the line 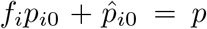 i.e. 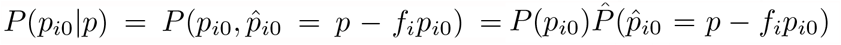 where 0 ≤ *p*_*i*0_ ≤ min{*p/f*_*i*_, 1} and 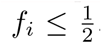. It follows that 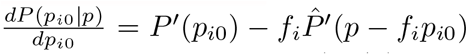, where the derivatives *P*′ and 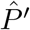 are negative. The distribution *P* (*p*_*i*0_|*p*) must therefore have a mode at *p*_*i*0_ = 0 or *p*_*i*0_ = min {*p/f*_*i*_, 1} or both, but nowhere else. The variance 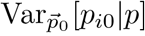 will be greatest in the two-mode case, so we assume a single mode at *p*_*i*0_ = 0 without loss of generality i.e. strictly decreasing *P* (*p*_*i*0_|*p*) with increasing *p*_*i*0_. Then, as *p* increases 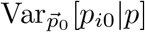 grows at rate approximately proportional to *p*^2^, because the most probable values lie near *p*_*i*0_ = 0 with an associated square deviation of (*p*_*i*0_ − *p*)^2^ = *p*^2^. Thus, when subpopulatoin allele frequencies are independent, 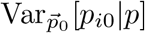 increases much faster than binomial ∼ *p*(1 − *p*) as 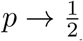, in accord with what we would expect from the frequency dependence of *G*_*ST*_ [5].

Allele frequency correlations among subpopulations can be caused by shared history or migration. In the case where isolated subpopulations descend from a shared ancestral population, allele frequencies will initially diverge binomially, tending to the independent case above as correlations decay due to drift.

The effects of migration among subpopulations are more complex, since sub-populations no longer evolve independently, and there can be nontrivial migratory relationships between subpopulations due e.g. to spatial structure. Given this complexity and our primary focus on closed laboratory populations, we leave detailed analysis of this scenario for future work. We note, however, that migration acts to eliminate subpopulation differentiation, moving the evolutionary dynamics in the population closer to admixture. It would therefore be surprising if migration between subpopulations created non-binomial allele frequency divergence at the population level.

### S3 Fluctuating selection and temporal autoco-variances

In the main text we analyzed the among-locus variance in Δ_*t*_*p* = *p*_*t*_ − *p*_0_, the total allele frequency change over *t* generations. Here we discuss the relative contributions of persistent directional selection versus fluctuating selection to this variance, and the relationship of this variance to the temporal autocovariance studied by [6]. As in the main text, we follow a cohort with the same initial major allele frequency *p*_0_ (not written explicitly to simplify the equations).

The total allele frequency change after *t* generations can be decomposed into a sum over the intervening *t* generations generations 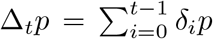, where *δ*_*i*_*p* = *p*_*i*+1_ − *p*_*i*_ (in the empirical studies considered in the main text, these intervening *δ*_*i*_*p* are not measured). From Eq. (2), we have *δ*_*i*_*p* = *s*_*i*_*p*_*i*_(1 − *p*_*i*_) where *s*_*i*_ (*i* = 0, …, *t* − 1) is the total selection coefficient in generation *i*.

The contribution of selection to allele frequency divergence in an unbiased (*E*[Δ_*t*_*p*]=0) allele cohort is given by the among-locus variance Var_*s*_(*E*[Δ_*t*_*p*|*s*_0_, …, *s*_*t* −1_]).Assuming that the *s*_*i*_ are all small and the total allele frequency change is not too large (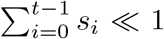 and | *s*_*i*_ | ≪ 1), the expected allele frequency change is 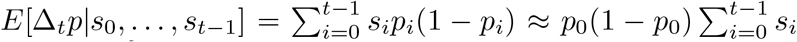 (dropping terms of order *s*^2^). This gives

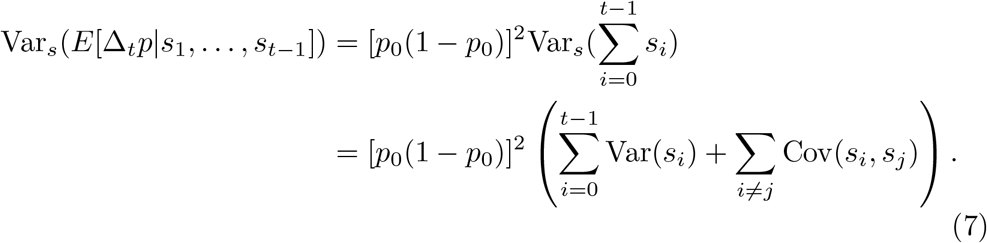

The two sums on the right can be interpreted respectively as: the divergence contribution from fitness variation within the intervening generations; and the divergence contribution arising from temporal consistency in that fitness variation across the intervening generations.

We assume that the total selection coefficients *s*_*i*_ are random variables with probability distribution that may depend on time *i* and locus *l*. The expectation of this distribution, denoted *E*[*s*_*i*_ | *l*], represents the consistent selective pressure in generation *i* at locus *l*. This consistent pressure could be constant over time, cycle periodically, or have no temporal pattern. The deviation of *s*_*i*_ from *E*[*s*_*i*_|*l*], denoted *ϵ*_*i*_ = *s*_*i*_ − *E*[*s*_*i*_ | *l*], represents a random selective component arising from unpredictable environmental factors or randomness in the genetic backgrounds experienced by each allele due to genetic drift, assortment and recombination. The variance of *ϵ*_*i*_, Var(*ϵ*_*i*_|*l*), measures the intensity of stochastic fluctuations in *s*_*i*_ at locus *l*.

First consider the simplest case where *t* = 1, such that only the within-generation divergence from generation *i* = 0 contributes to (7). Applying the law of total variance,

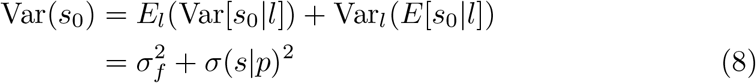

where we have used the fact that Var_*l*_(*E*[*s*_0_|*l*]) = *σ*(*s*|*p*)^2^, and introduced the notation 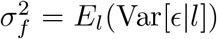 for the locus-averaged intensity of fluctuating selection.

Now consider the *t >* 1 case. For illustration we suppose that *s*_0_, …, *s*_*t−*1_ are independent and identically distributed (iid) random variables (but allowed to be different between loci). This represents a situation where there is no systematic time-dependence in the *s*_*i*_; the only time dependence is due to temporally-uncorrelated fluctuations. The within-generation contribution to divergence is then given by

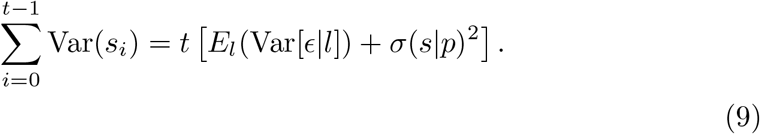

The within-generation divergence thus grows linearly with time analogous to a random walk. This is to be expected since the within-generation divergence represents the effects of among-locus fitness variation solely on a generation-by-generation basis regardless of how that variation is correlated over time. In the case where there is no systematic among-locus variation *σ*^2^(*s* | *p*) = 0, there remains a divergence due to fluctuating selection with a non-binomial frequency dependence [7].

On the other hand, the between-generation selective divergence is

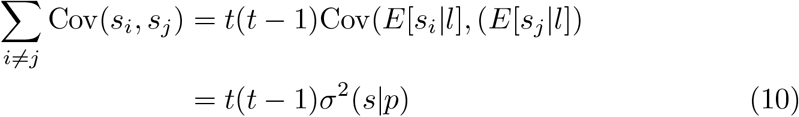

which grows quadratically over time. Sustained directional selection thus generates divergence an order of magnitude faster than fluctuating selection. Therefore, in the iid case, the overall divergence starts to be dominated by between-generation covariances for large *t* when we have *t*^2^ ≫ *t*.

The empirical studies in the main text have *t* ∼ 10, which is large enough that the covariance contribution would dominate for iid *s*_*i*_. However, the iid assumption is often not realistic. For neutral alleles, the covariance between *s*_*i*_ and *s*_*j*_ is expected to decay exponentially with increasing time separation | *j* − *i* |. In the two-locus case where the neutral allele is hitchhiking with one selected allele, linkage disequilibrium (and thus covariance) decays at rate ∼ (1− *r*)^|*j−i*|^ [8] where *r* is the recombination rate. The multilocus case similarly involves exponential decay averaged over all linked sites under selection [9]. Nevertheless, even if recombination destroys linkage disequilibrium so rapidly that only concurrent generations | *i* − *j* | = 1 covary, there are still *t* − 1 such pairs contributing to (7). Thus, the among-locus allele frequency divergence may contain a substantial contribution from the accumulation of temporally consistent directional selection, represented by the among-locus temporal autocovariances Cov(*s*_*i*_, *s*_*j*_).

### S4 Selective perturbation of the drift variance

Here we evaluate the influence of selection on the drift contribution to allele frequency divergence, *E*_*s*_[Var(Δ_*t*_*p*|*p*, {*s*_0_, …, *s*_*t−*1_})]. We do so by deriving a recursion for Var(Δ_*t*_*p*|*p*, {*s*_0_, …, *s*_*t−*1_}) = Var(*p*_*t*_|*p*_0_, {*s*_0_, …, *s*_*t−*1_}) going backwards along the allele frequency trajectory *p*_*t*_, …, *p*_1_, *p*_0_ where *p*_0_ = *p*.

We start with the neutral case *s*_0_ = … = *s*_*t−*1_ = 0. Applying the law of total variance to account for the variance added to *p*_*t*_ due to the allele frequency variance from the preceding generation *p*_*t−*1_, we have

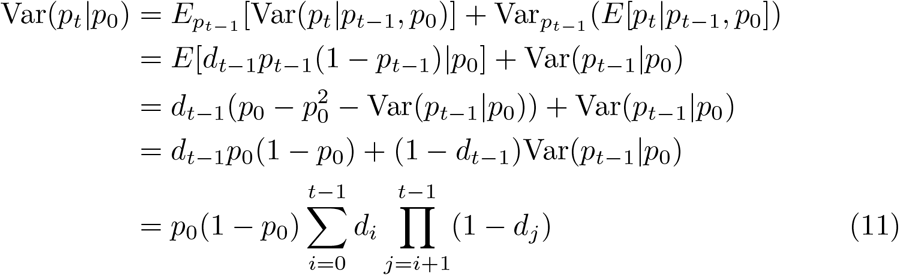

where *d*_*t*−1_ is the variance coefficient for Var(*p*_*t*_ |*p*_*t*−1_) and the last line is obtained by recursion. The variance coefficient for Var(*p*_*t*_|*p*_0_) is thus 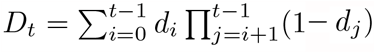.

We now evaluate how selection changes Eq. (11). We only evaluate the first order effects of selection since we focus on the scenario where selective allele frequency changes are small over the interval *t* (i.e. 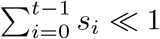 and |*s*_*i*_| ≪ 1).

First, selection perturbs the variance over one generation such that Var(*p*_*i*_|*p*_*i−*1_, *p*_0_) = *d*_*i−*1_(*p*_*i−*1_(1 − *p*_*i−*1_) + *δ*_*i−*1_) where the perturbation *δ*_*i−*1_ is of order *s*_*i−*1_ (for brevity we do not explicitly write the conditional dependence on {*s*_0_, …, *s*_*t−*1_}).

Second, we now have (to first order in *s*)

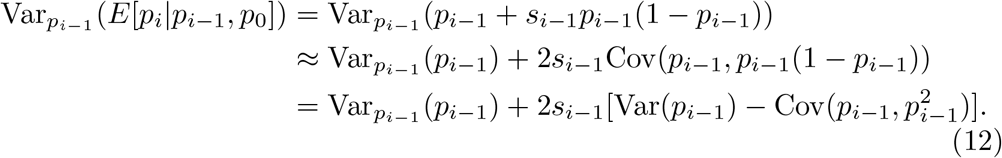

The remaining covariance can be written as

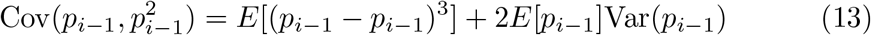

We assume that the initial cohort allele frequency *p*_0_ is sufficiently far from fixation, and the duration of divergence *t* is small enough, that *p*_*i*_ is close to symmetrically distributed for all 1 ≤ *i* ≤ *t* − 1, such that the skewness term *E*[(*p*_*i−*1_ − *p*_*i−*1_)^3^] can be neglected. Moreover, the selective change to *E*[*p*_*i−*1_] is *O*(*s*^2^) since the covariance is multiplied by *s*_*i−*1_. We thus obtain,

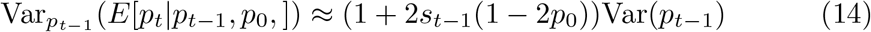

to first order in *s*. Consequently, the neutral result Eq. (11) becomes

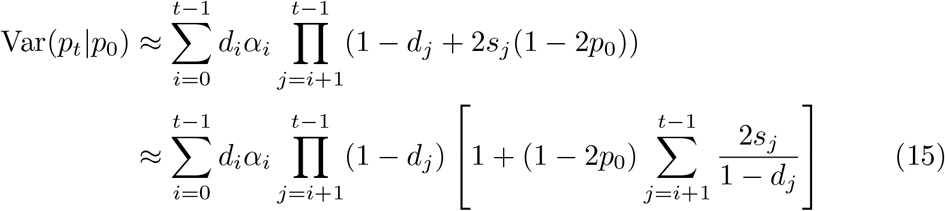

where *α*_*i*_ = *p*_*i*_(1 − *p*_*i*_) + *δ*_*i*_.

Taking the expectation *E*_*s*_ and substituting 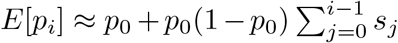, which implies 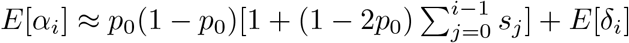, we obtain

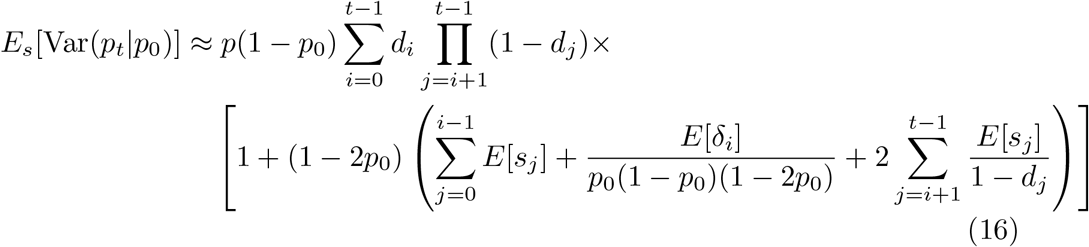

to first order in *s*.

Observe that the term in large parentheses in Eq. (16) is closely related to the expected selection coefficient for the accumulated allele frequency change after *t* generations, 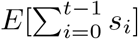. One complication is the presence of the within-generation perturbation *δ*_*i*_, the exact form of which will in general depend on population dynamic specifics. However, the contribution from the term containing *δ*_*i*_ will not be important when *t* ≫ 1, as is the case in the present study (*t* ∼10 generations). Moreover, note that in the Wright-Fisher model, selection is usually incorporated by applying the selective perturbation to gamete frequencies prior to binomial sampling; this gives Var(*p*_*i*+1_|*p*_*i*_, *p*_0_) = *d*_*i*_*p*_*i*+1_(1 − *p*_*i*+1_) and thus *δ*_*i*_ = *p*_*i*_(1 −*p*_*i*_)(1 −2*p*_*i*_)*s*_*i*_; hence *E*[*δ*_*i*_]*/*[*p*_0_(1 −*p*_0_)(1 −2*p*_0_)] ≈*s*_*i*_. Similar behavior occurs in the continuous-time Moran model (see Appendix in [10]). Thus, the *δ*_*i*_ term could plausibly be proportional to *s*_*i*_ (with no frequency dependence) in many populations.

Another complication is the extra weight given to later generations due to the doubling factor in the term 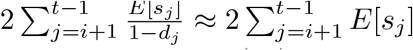. This weighting means that the term in large parentheses in Eq. (16) will have a value close to 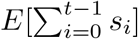. That is,

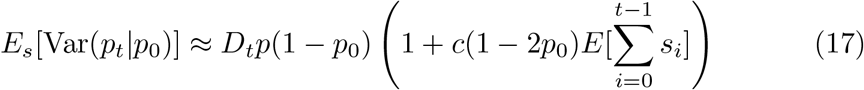

where *c* is a constant factor of order 1.

### S5 Measurement error

In this section we show that the major sources of allele frequency measurement error do not systematically alter the frequency independence of *C*_*t*_ in main text Eq. (1). Population allele frequencies are denoted *p* while estimated allele frequencies are denoted 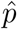.

Suppose that the measurement variance satisfies 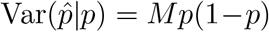 where *M* is frequency independent and can also differ between generations. Applying the law of total variance (for any two random variables *X* and *Y*, Var(*Y*) = *E*_*X*_ [Var_*Y*_ (*Y* |*X*)] + Var_*Y*_ (*E*_*X*_ [*Y* |*X*])), alleles with the same initial frequency *p*_0_ will have variance,

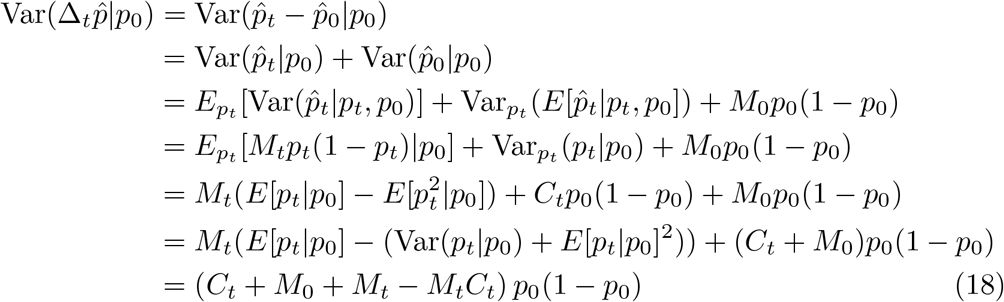

where we have used the fact that *E*[*p*_*t*_ |*p*_0_] = *p*_0_ for neutral alleles. Therefore, measurement error does not change the frequency dependence of neutral cohort variances, provided that our assumption of frequency independent *M*_*t*_ is true. We now show the latter accounting for different sources of error. Following [11], we divide measurement error into three main sources that will be considered in turn: population sampling, pooling and amplification of the sampled DNA, and finite read depth.

#### Population sampling

Random sampling of individuals generates a binomial sampling error, such that the sample allele frequency *p*_*S*_ has variance Var(*p*_*S*_ |*p*) = *p*(1 −*p*)*/n*_*S*_ where *n*_*S*_ is the sample size. Using the law of total variance, the contribution of this sampling error is

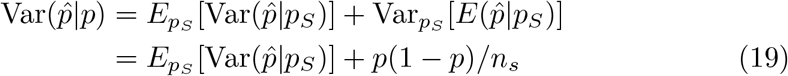

The term 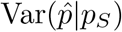 represents the error in 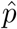 introduced after population sampling, which we now evaluate.

#### Read sampling

Let *r* be the number of reads of the reference allele *A* recorded by the sequencer at a locus with a total of *r*_*T*_ reads. The allele frequency estimate of *A* is 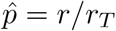. We initially restrict our attention to loci that have the same read depth *r*_*T*_ ; below we incorporate the effects of among-locus variation in *r*_*T*_. Reads are assumed to be generated by random sampling with replacement from the DNA pool. Reference allele reads are thus binomially distributed *r* ∼Bin(*r*_*T*_, *p*_*P*_) where *p*_*P*_ is the allele frequency of *A* in the sequenced DNA pool.

If all allele copies sampled from the population contribute equally to the sequenced DNA pool, then *p*_*P*_ = *p*_*S*_. Errors introduced during pooling and amplification cause unequal contributions and thus variance in *p*_*P*_ among sequenced loci. We assume that pooling+amplification error do not systematically bias allele frequencies such that 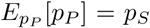. Applying the law total variance to 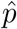 for fixed *r*_*T*_ gives

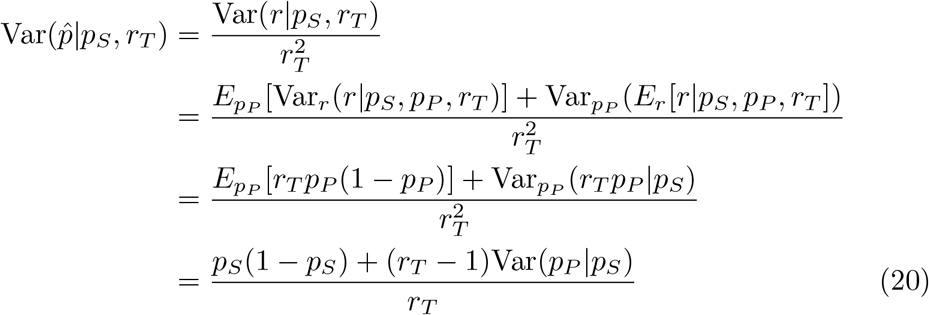

where we have used the identity *E*[*X*^2^] = Var[*X*] + *E*[*X*]^2^ in the last line. Note that the conditionality on *r*_*T*_ has been dropped in 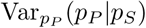 because the DNA pool frequency *p*_*P*_ is independent of *r*_*T*_. We now incorporate the effects of pooling and amplification to show that 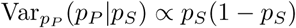.

#### Pooling+amplification

Let *F*_*i*_ denote the proportion of the DNA pool at the focal locus originating from the *i*’th sampled allele copy, where *i* = 1, …, 2*n*_*S*_ for diploids. In the case of equal contributions from each sampled allele copy we simply have *F*_*i*_ = 1*/*2*n*_*S*_, but in general the *F*_*i*_ are random variables with unknown probability distributions denoted *P* (*F*_*i*_) (ref. [11] assumes a Dirichlet distribution for the *F*_*i*_). We have 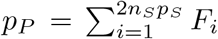 where 2*n*_*S*_*p*_*S*_ is the number of sampled allele copies containing the focal allele (i.e. we order the haplotype labels *i* such that the reference allele copies appear first). Since the *F*_*i*_ sum to one, we have analogous to (1),

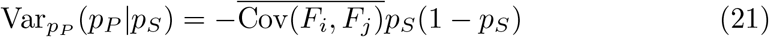

where 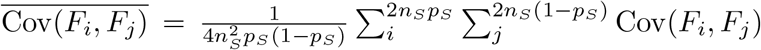 is the mean covariance of the DNA pool contributions between the focal and non-focal alleles.

Similar to exchangeability in the Cannings model, pooling and amplification are blind to allele copy identity and should therefore affect allele copy probabilities symmetrically (see S1). This implies 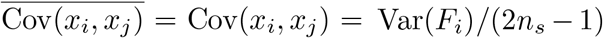 independent of *i* and *p*_*S*_. Combining the this result with (20) we therefore obtain

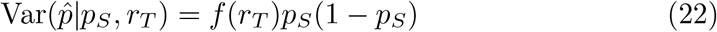

where *f* (*r*_*T*_) is independent of *p*_*S*_.

#### Variable read depth

Read depth varies across loci according to some probability distribution *P* (*r*_*T*_). This variability in incorporated by integrating out the dependence on *r*_*T*_

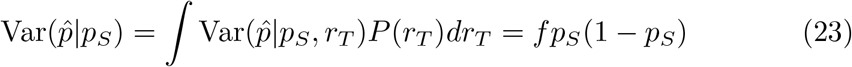

where *f* = ∫*f* (*r*_*T*_)*dr*_*T*_. Substituting this result into (19) we finally obtain

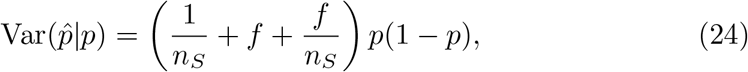

where the term multiplying *p*(1 −*p*) is frequency-independent as claimed at the start of S2.

**Figure 1:**
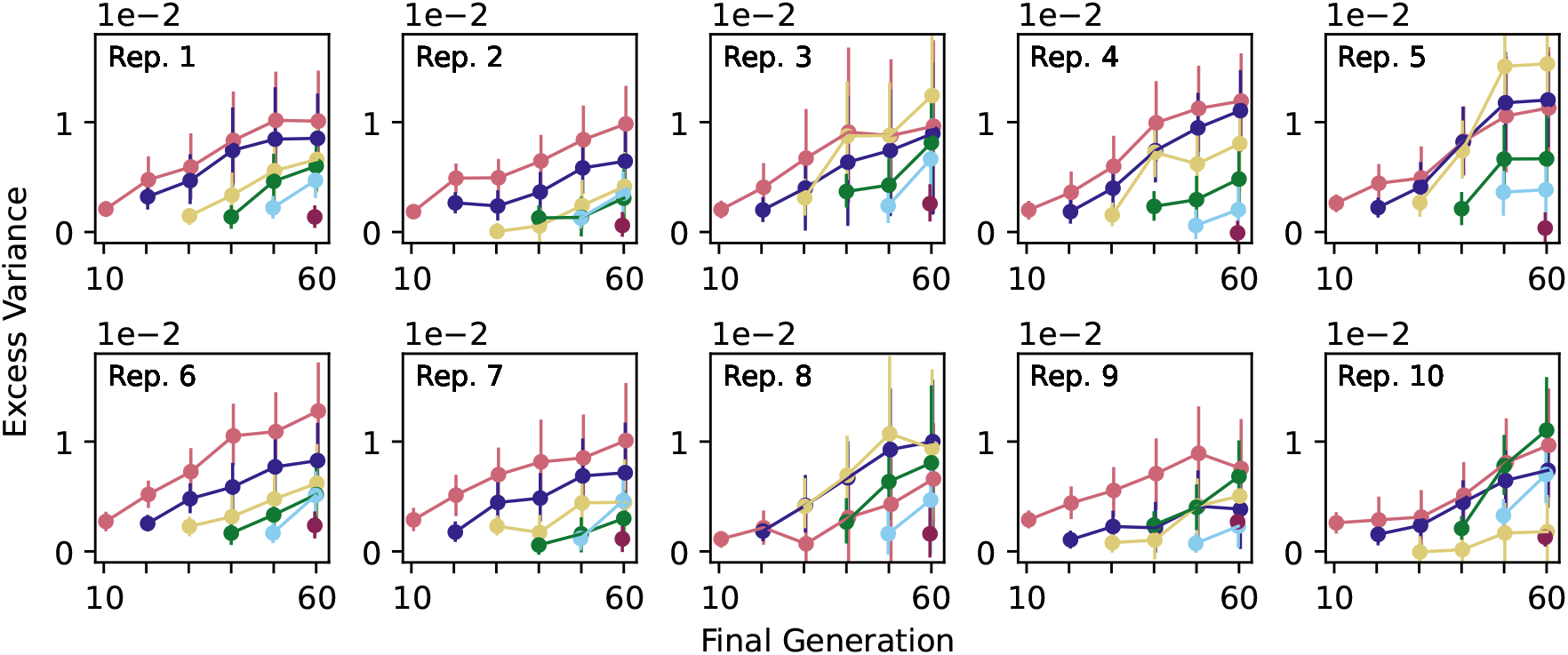
Identical to Fig. 2A in the main text but including all 10 replicates from Barghi et al.

**Figure 2:**
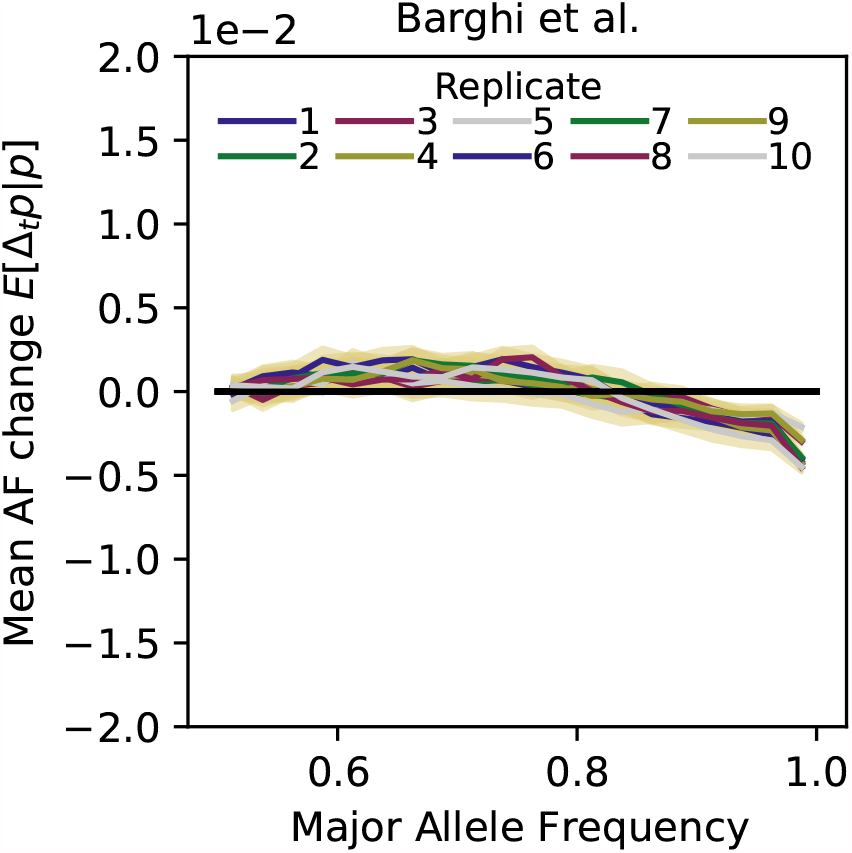
Barghi et al. expected Δ_*t*_*p*, first iteration, all replicates. 95% 1MB block bootstrap confidence intervals are too narrow to be easily visible.

**Figure 3:**
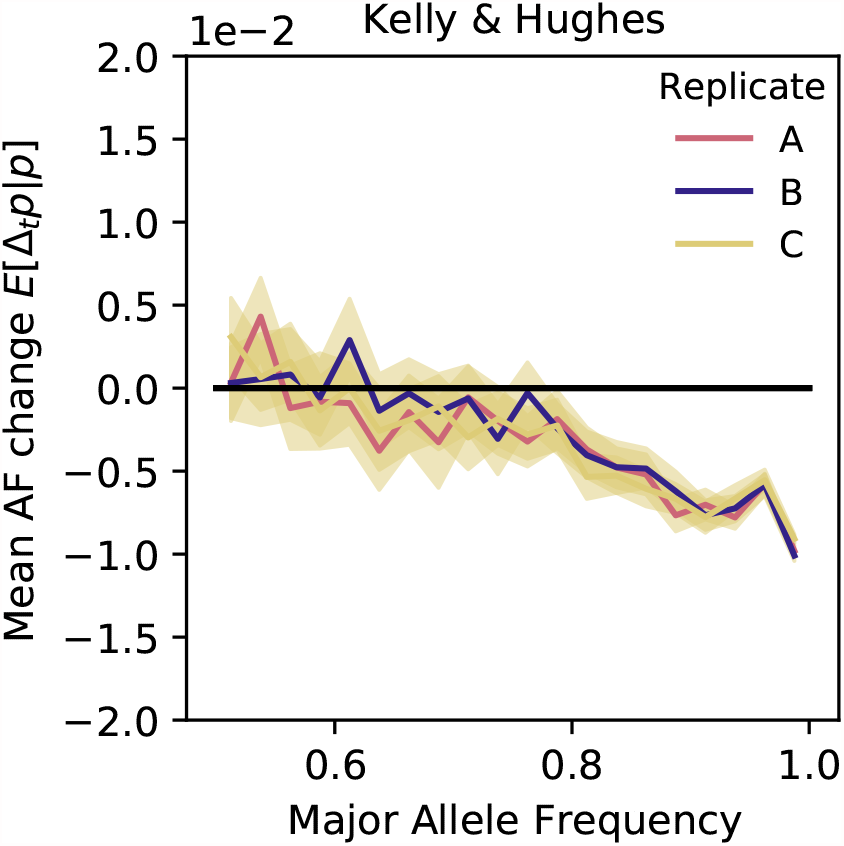
Kelly et al. expected Δ_*t*_*p*, all replicates, with 1MB Block bootstrap 95% confidence intervals.

**Figure 4:**
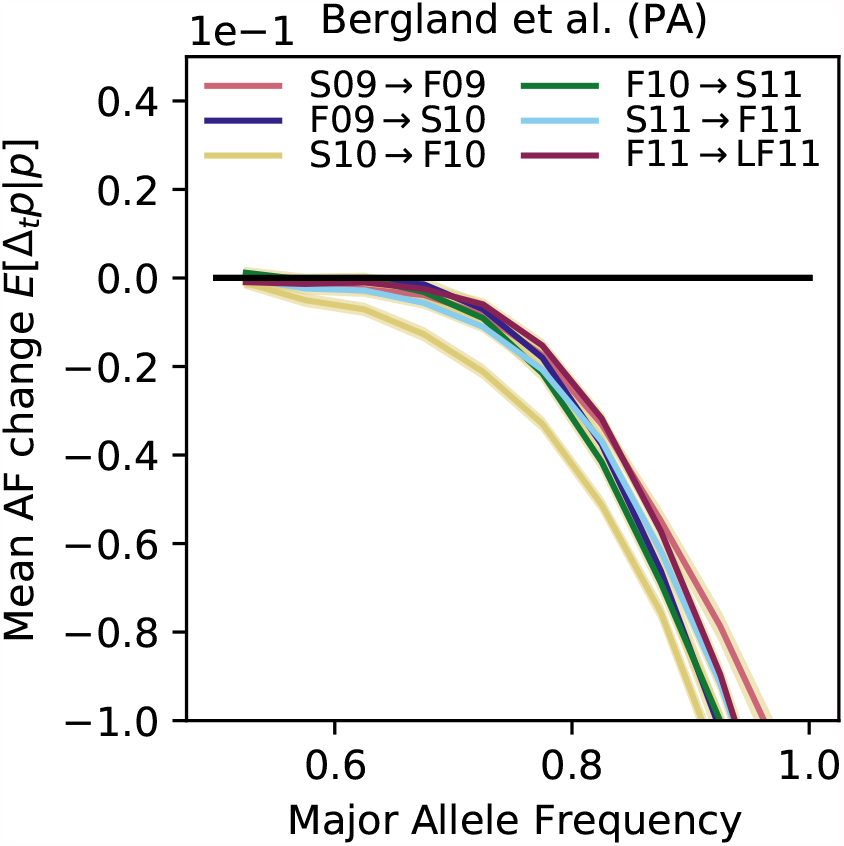
Bergland et al. expected Δ_*t*_*p* over all seasonal iterations. 95% 1MB block bootstrap confidence intervals are too narrow to be easily visible.

